# Foraging fidelity and individual specialisation in a temperate bat Myotis Nattereri

**DOI:** 10.1101/713750

**Authors:** Simone Mordue, Aileen Mill, Mark Shirley, James Aegerter

## Abstract

1. Bat populations have declined globally over the last century largely due to anthropogenic change. Many temperate forest species of bat appear loyal to their foraging sites however, conservation of these sites rather than just habitat types is rarely considered and is essential to protect bat populations. It is not clear whether site fidelity in bats is species-specific or a more general trait or why it is exhibited but behaviour patterns could be important for conservation and management objectives. Foraging variation may occur due to ‘individual specialisation’, such that individuals differ significantly in their prey or habitat utilisation, independent of class-effects. If bats do exhibit individuality in their habitat choice, then protection of a mosaic of habitats rather than single preferred habitat per species may be critical to their conservation.
2. The goal here was to determine whether Natterer’s bats show fidelity in their foraging choices and whether they show individual specialisation in their foraging habits.
3. Thirty-four individual bats were tracked for at least one full night, from two different sites.
4. Site fidelity in Natterer’s was consistent across a range of intervals (months and years) despite contrasting seasonal contexts. Individuals repeatedly exploited specific foraging locations and showed individual specialisation in their habitat use which is consistent with the behaviour of a territorial species.
5. Studies designed to inform conservation and management of temperate bats should attempt to maximize the number of individuals from which movement data is sought, whilst ensuring that data represent a coherent and meaningful measure of behaviour such as a single full night. Bat conservation may need to shift from general descriptions of habitat preferences to considering individual specialisation in habitat use. Designing conservation strategies resilient to environmental change might then advocate protecting a mosaic of habitats to preserve the habitat specialisms of many individuals and enhance their productivity rather than advocating the preservation of a single preferred habitat only suited to a few individuals.

## Introduction

Globally bat populations have declined considerably over the last century largely due to habitat loss, hunting, or disease (1) with recovery predicted to be slow (2). Determining habitat types required for effective conservation of bats can be difficult compared to terrestrial mammals of similar size, due to their ability to fly, which enables them to travel long distances, allowing use of a wide range of resources across a wider variety of habitats. In summer bat roosts and foraging sites can be kilometres apart permitting them to assemble widely dispersed resources they require for survival and reproduction from across extensive landscapes (3).

Flight and the large areas it allows bats to exploit, can also expose individuals to a range of anthropogenic threats such as development (inc. residential, commercial and infrastructure) or sources of mortality such as wind-turbines (4–9) and roads (10–13) on a nightly basis. These threats can also affect roosts, reduce or degrade available foraging habitat or interfere with the connectivity between habitats (1). Although the conservation of roost sites is important for the maintenance of bat populations e.g. (14), roost provision or the mitigation of roost loss is already common in the management of bat populations. In addition to this, many species use a network of alternative roosts and cope with the loss of a few (15–18). Conversely, bats appear loyal to their foraging sites despite roost loss (17). Hence, the conservation of foraging sites (rather than just habitats) may be as important as roost sites in the conservation of the species, although rarely considered. Further, it is not clear that there are tools to confidently identify the most important foraging areas in the landscape (rather than particular preferred habitat types). This is of some concern especially as determining priorities for policy led management of habitats requires a robust understanding.

It is not clear whether site fidelity in bats (14, 19) is specific to temperate forest bats, a more general behaviour found in other species or why it is exhibited. Initial work by Egert-Berg et al (20), suggests that many bats may demonstrate a sufficient knowledge of their foraging landscape to choose to repeatedly return to specific productive sites. Male Noctule bats *Nyctalus noctula* (an aerial hawking species), also show repeated use of the same foraging strategy (foraging trajectories and areas) (21).

Foraging animals are expected to make choices in order to minimise energy expenditure whilst maximising energy intake, therefore site fidelity may be a way of saving energy in the decision of which habitat to use next (22) e.g. a previously beneficial habitat may have a higher probability of providing sufficient resources than an unexplored new habitat (23). More recently the idea of bats exhibiting site fidelity has been extended to suggest the territorial defence of feeding areas through the use of social calls (24, 25). Territorial defence of food resources is thought to minimize feeding competition (26), maximize feeding efficiency through familiarity with the distribution of food resources (27, 28) or directly impact reproductive success (29). Site fidelity or territoriality could have significant impacts on conservation and management objectives as well as the tools and measures used to describe bats use of space.

Studies to support bat conservation may need to shift from general descriptions of habitat preferences and the assumptions that as long as these are accessible, Favourable Conservation Status can be maintained at specific sites, to considering other aspects of behaviour. Bats like many other mammals may have individual preferences (30, 31) and this may interact with the emerging study of bat sociality, such that dominant females have preferential access to the best resources (32), maternal inheritance of foraging sites occurs, or bats exhibit territoriality or personality (33) which affects their access to foraging areas. Sexual segregation may also occur with females only utilising more favourable habitats to the exclusion of males who may be found more abundantly in marginal habitats (34). Alternatively, foraging variation may occur due to ‘individual specialisation’, such that individuals differ significantly in their prey or habitat utilisation, independent of class-effects (35).

Individual specialisation has important evolutionary (35), ecological and management implications as interactions between individuals and environment are not uniform across a landscape (36). If bats do exhibit individuality in their habitat choice, then there may no preferred habitat per species as often asserted by the literature e.g. (37–40) but instead a mosaic of habitats may be critical to their conservation in a changing environment. Therefore, the influence individual expressions of behaviour may have on our understanding of species requirements should be accounted for in measures of resource use and considered in conservation planning and management.

Here the aim is to produce robust descriptions of where individual bats forage and the habitats they use. It is also to explore the fidelity individual bats show in their foraging choices and the possibility that Natterer’s bats in this study show individual specialisation in their foraging habits. The consequences of the observed behaviour and the implications for conservation management are then discussed.

## Materials and methods

Natterer’s bats from two sites in Northern England were radio tracked and continuous contact with each bat for full nightly foraging trips was achieved, from dusk to dawn. Foraging site fidelity of individual bats was measured by the degree of spatial overlap of data collected at varying intervals (days, months and years) to determine whether the data collected over a single night has the potential to represent a more substantive and general description of individual behaviour.

### Study sites

Natterer’s bats *(Myotis nattereri)* were caught at a roost in a church at Low Catton, East Yorkshire, UK (53.98° N, 0.93° W: altitude 15m) between May and August in 2003 and from woodlands on the Wallington Estate, Northumberland, UK (55.15° N, 1.96° W: altitude 160-200m) between May-September of 2013, 2014 and 2015 (Supplementary 1). Low Catton is a very small rural village in a mixed agricultural landscape typical of lowland England (mainly arable though with some pasture and small scattered parcels of woodland). Wallington Estate is a patchwork of parkland, lakes and woodland, within a mixed pastoral landscape including arable and woodland components as well as open moorland; typical of an upland agricultural landscape in England.

### Bat capture and radio-tracking

Bats were captured on an approximately weekly schedule at roosts using a static hand net or harp trap attached to an extendable pole. The frequent roost switching behaviour of the bats at Wallington also required their capture from free flight, using mist nets or harp traps, and an acoustic lure. All disturbances at roosts, as well as the capture, handling and marking of bats were carried out under licence from Natural England (2014-6454-SCI-SCI). and a full ethical assessment was carried out and approved by the Animal Welfare Ethical Review Board (AWERB). Captured bats were described noting; sex, age (adult/juvenile), reproductive condition (pregnant or lactating; by palpation of the abdomen), forearm length (0.1mm) weight (to 0.1g) and any existing mark. Unmarked bats were marked with a unique ring (2.9mm Alloy; BCT, England). Selected bats were fitted with radio transmitters (Pip AG317; Biotrack, Essex, England) attached to the skin between the scapulae using a hypoallergenic dermal adhesive. A small patch of fur was trimmed at the point of attachment to ensure reliable adhesion. A maximum of two bats were marked with transmitters in one tracking session to ensure that a complete and continuous night of data could be collected from all deployed tags within the short duration of their batteries (7-10days). Forecasts of inclement weather were considered to ensure the timing of tag deployment could always yield robust data.

Individual bats were usually radio-tracked by single workers using the close approach method (41) using a Telonics TR-4 receiver (Telonics, Arizona, USA) attached to three-element flexible Yagi antenna or vehicle mounted omni-directional antennae (42). Bats were tracked from their emergence from roosts until their return, with their locations recorded at 10 minute intervals. 10 minute intervals were selected to prevent temporal correlation between consecutive fixes whilst still recording regular movement patterns (42).

Due to the difficulty in obtaining triangulations from fast flying animals (43) especially across undulating terrain such as that of our field site at Wallington, approximate locations of bats were inferred using the strength, direction and variability of signal and recorded onto a large-scale map as polygonal observations. Polygons described an area containing the bat’s true location later used in determining their foraging cores. The more traditional description of triangulated points (which uses only the bearing) and subjective estimation of error associated with these were both impractical (co-ordination of observers in difficult terrain) and sacrificed information and tracking effort (triangulation requires paired trackers) without necessarily being more objective. Tracking was undertaken in two phases. In the first, priority was given to simply maintaining contact with the bat and establishing its general foraging strategy (i.e. commuting routes, approximate location of its favoured foraging patches and the schedule of behaviours); trackers commonly stayed closer to access routes and vehicles to ensure a rapid response to unexpected bat movement; though this often resulted in more uncertain estimates of location. In the second phase, once it was felt that the general strategy used by that animal was known, trackers planned to optimise the quality of data by anticipating bat behaviour, and committing themselves to closer approaches on foot where this was possible. However, the intention was always to ensure continual contact with the bat throughout its period of activity. Data was only collected from tracking in the second phase of work for every bat and a complete night of data was only considered to have been recorded if continual contact with the bat was maintained and the night was not unexpectedly interrupted by inclement weather (not uncommon at these locations). For most bats a single complete night of data was collected during the second phase, although some were re-tracked at varying intervals either within a summer season or between years. The minimum time between tracking periods was one night. Phase 1 tracking often took 2-3 nights (for bats travelling long-distances quickly or those traversing difficult to cross barriers in the landscape such as rivers), and at least one additional night of effort to secure a complete, continuous and uninterrupted night of foraging data. A number of different trackers were used to establish general foraging strategies (phase 1 tracking), especially where a number of bats were tagged simultaneously. However, the same tracker (SM at Wallington, JA at Low Catton) undertook all data collection during phase 2 to ensure a consistency in the inference of location and the confidence of its recording. Nights of tracking data with continuous gaps of more than 20 minutes were excluded from the analysis. All roost positions were recorded using a handheld GPS device. Observations were digitised using ArcGIS (v.10.2; ESRI). Subsequent analysis was carried out in R (v. 2.1).

Core foraging patches representing areas of high use were described for each complete bat-night and used to inform spatial analyses. Observations were transformed into clouds of points by placing a single spatially randomised point into each polygonal description of location. Cores were then identified using a non-parametric clustering approach (clusthr function in adehabitatHR: 95% inclusion). This was repeated 5 times and the resulting cores were then intersected to find the areas common to all 5 iterations by sorting and overlaying each (gIntersection function in rgeos). These core patches used by bats on any given night represent locations that include the centres of high-density use and the area most exploited for foraging. They also represent the statistical unit in all subsequent analyses.

### Data analysis

#### Exploring habitat use

Core patches for each individual from the first full night of tracking data collected, were intersected with a land cover layer, LCM2007; (44) to describe the composition of the habitats most used by the bats. To quantify individual specialization using proportion data, Roughgarden’s index (45) was used which compares within-individual components of niche width (WIC) to the total niche width exhibited by a population (TNW). Calculations of WIC and TNW were carried out in R (RinSp using the ‘PSicalc’ function(46)). For each individual, the proportional similarity index (PS_i_) was calculated following Fodrie et al. (47) based on habitat deviations in an individual’s habitat use relative to population level, average habitat use (approaching 1 = more generalist; approaching 0 = more specialised). Mean PS_i_ among individuals was used to determine the average amount of specialisation in habitat use across all bats in this study and individuals were deemed to be specialists if their PS_i_ value was below the mean population Ps_i_ value. Monte Carlo permutations were run with 999 replicates to test whether observed WIC/TNW and PS_i_ values differed significantly from a random distribution of values subsampled from the population.

#### Foraging fidelity

The potential of a single complete night of radio-tracking data to act as a proxy for the long-term description of an individual’s foraging strategy was explored. Pairwise comparisons of foraging area metrics were carried out for all bats including those tracked only once and those tracked repeatedly, either within the calendar year, or between years to describe the similarity of foraging strategies across differing intervals. For each bat the core area was compared between nights and the proportion of area overlap was calculated for each (using the gIntersection function in the rgeos package (48)). This was compared to the proportion of overlap the core area had with every other bat core foraging area. Multiple-response permutation procedure (function MRPP, in the vegan package (49)) analyses were conducted using the Euclidian distance metric and 1,000 iterations with individual proportion overlap as the response variable. To explore whether individual foraging strategies/behaviour measured at varying intervals was repeatable, foraging distance (from roost to the most heavily used core) was compared to foraging schedule (time from emergence to arrival at the most heavily used core) across all of the data with observations of the same bat compared to observations of different bats (using the rpt function in package rptR (50), with 1000 permutations) with individual distance and foraging schedules as the response variables. To determine whether individuals had similar foraging strategies and adhered to the optimal foraging theory, the relationship between distance travelled to the most used foraging patch was assessed along with the speed at which they travelled there using a linear model (lm) with Distance as the response variable and Speed and Site (Wallington/Low Catton) as the predictors.

## Results

Thirty-four individual bats, 17 from Wallington and 17 from Low Catton, were tracked for at least one full night (Additional file 1). Six bats were tracked over repeated years and 24 bats were tracked twice during the same year (Table 1). This produced 29 nights of data available for pairwise comparisons of foraging site fidelity at Wallington and 32 nights at Low Catton. There was an average of 32 observations per night (range 16-46) at an average of 20 different foraging patches (range 8-44) resulting in a mean foraging period of 5.3 hours (tracking usually represented most, if not all, of the period of dark at 55° N in summer (Table 2)). At Wallington the majority of bats foraged at more than one patch (9/17) although the proportion was slightly lower at Low Catton (6/17). As all 10 minute observations could describe foraging activity we assume for the purposes of this study that all observations carry the same weight.

**Table 1.** Number of individual bats tracked by site and year used to estimate foraging habitat overlap at two temporal scales, within-year (multiple foraging trips by one bat in one year) and between-year (one bat over multiple years).

| Site | Year | Bats tracked |  |
| --- | --- | --- | --- |
|  |  | Fidelity level |  |
|  |  | Within-year | Between-year |
| Low Catton | 2003 | 17 | No |
| Wallington | 2013 | 4 | Yes (4) |
|  | 2014 | 2 | Yes (4) |
|  | 2015 | 1 | Yes (2) |

**Between-year fidelity (Yes or No) indicates for which years multi-year animals were tracked with the number of individuals in each of those years in brackets.**

**Table 2.** Mean number of Radio tracking observations, patches and foraging time of Natterer’s bats tracked at Low Catton and Wallington

|  | Site | Mean | Range |
| --- | --- | --- | --- |
| Observations | Low Catton | 30 | 16-46 |
|  | Wallington | 34 | 25-41 |
| Patches | Low Catton | 25 | 12-44 |
|  | Wallington | 14 | 8-23 |
| Foraging time<br>(hours) | Low Catton | 5.0 | 2.66-7.66 |
|  | Wallington | 5.66 | 4.16-6.83 |

In terms of movement dynamics, the relationship between the distance of the roost of departure and the most used patch and the speed between the two, were of interest as they represent independent choices (Figure 1). Some bats choose to travel long distances and some bats choose to travel quickly but there was not a consistent relationship between the two (R^2^ = 0.217, F_(3, 30)_=4.062, p=0.95).

**Figure 1.**
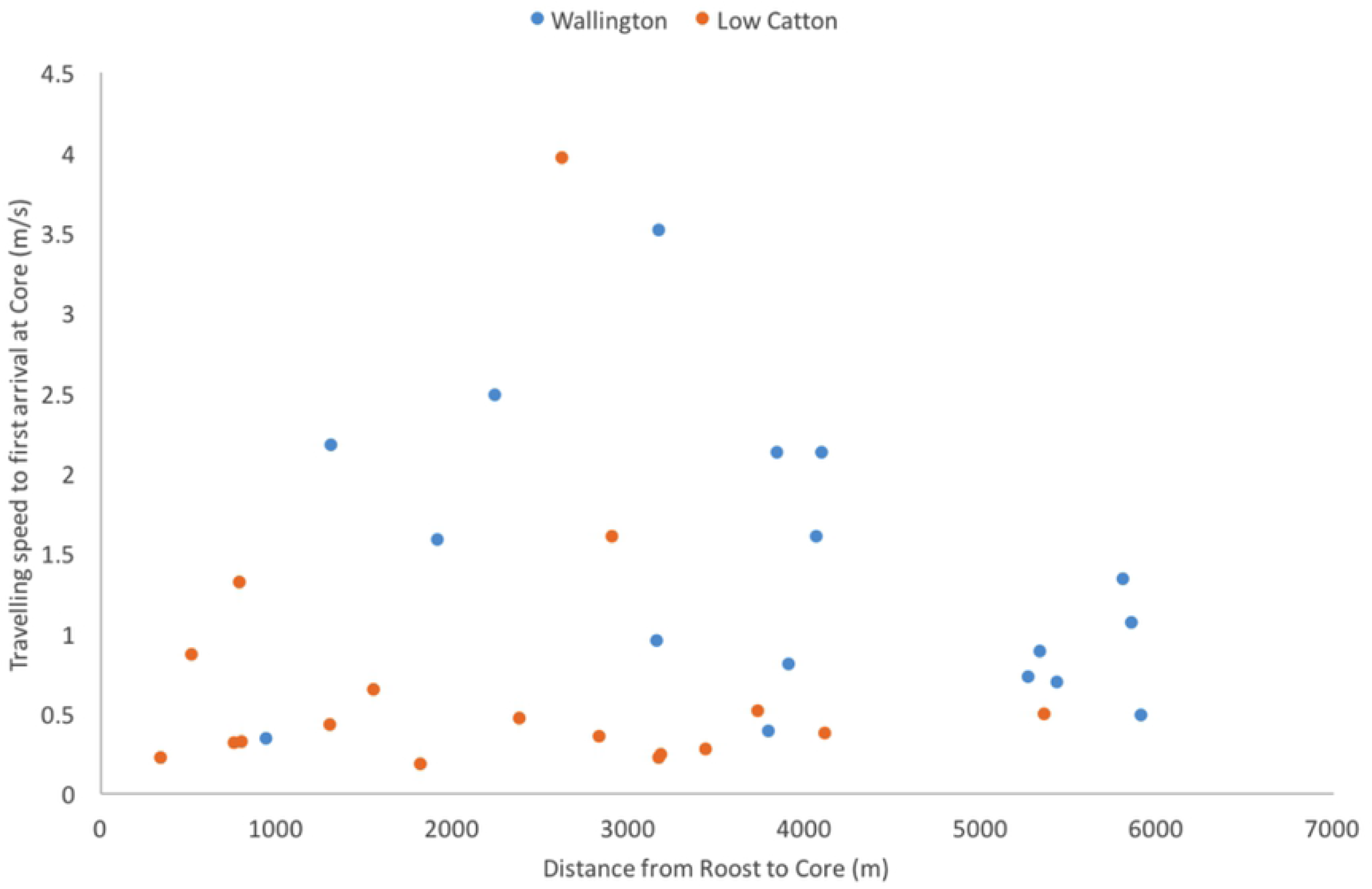
Distance from roost to core foraging area and speed for bats tracked at Wallington and Low Catton

### Foraging site fidelity

At both Wallington and Low Catton, a greater degree of foraging site overlap was observed for individuals tracked multiple times in the same year than between different bats (Figure 2,Figure 4,Figure 5; post hoc Tukey tests P<0.01; Low Catton same bat mean 0.92 ± 0.02(range 0.66-1); Wallington same bat mean 0.92 ± 0.06 (range 0.57-0.9); Low Catton different bats mean 0.29 ± 0.01 (range 0-1); Wallington different bats mean 0.62 ± 0.02 (range 0.25-0.99)).

**Figure 2.**
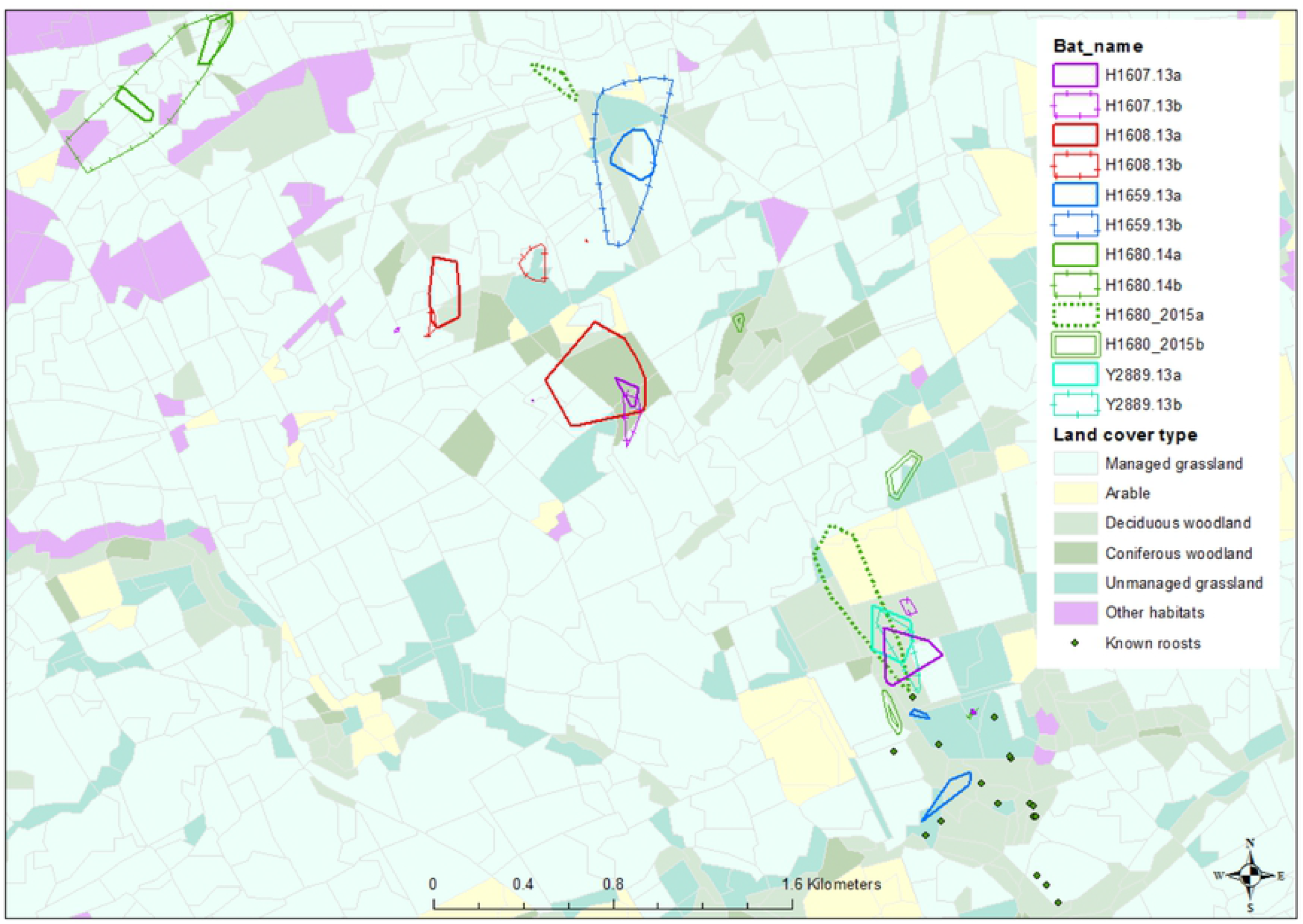
Foraging cores of bats (n=5) tracked repeatedly at Wallington within the same years (2013-2015) with 2007 Land cover data and roost locations

**Figure 3.**
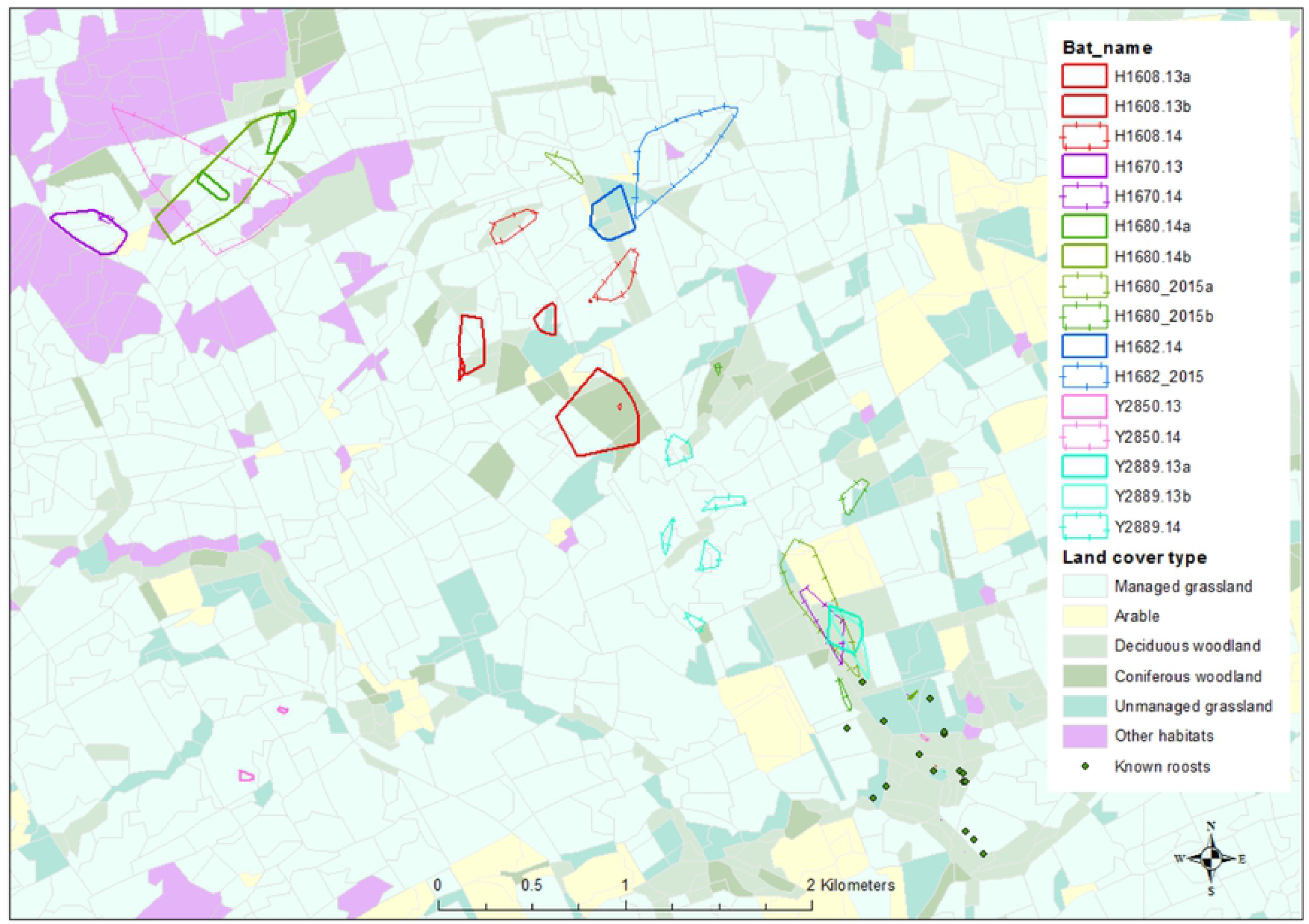
Foraging cores of bats tracked repeatedly at Wallington (n=6) between years (2013–2015) with 2007 Land cover data and roost locations

**Figure 4.**
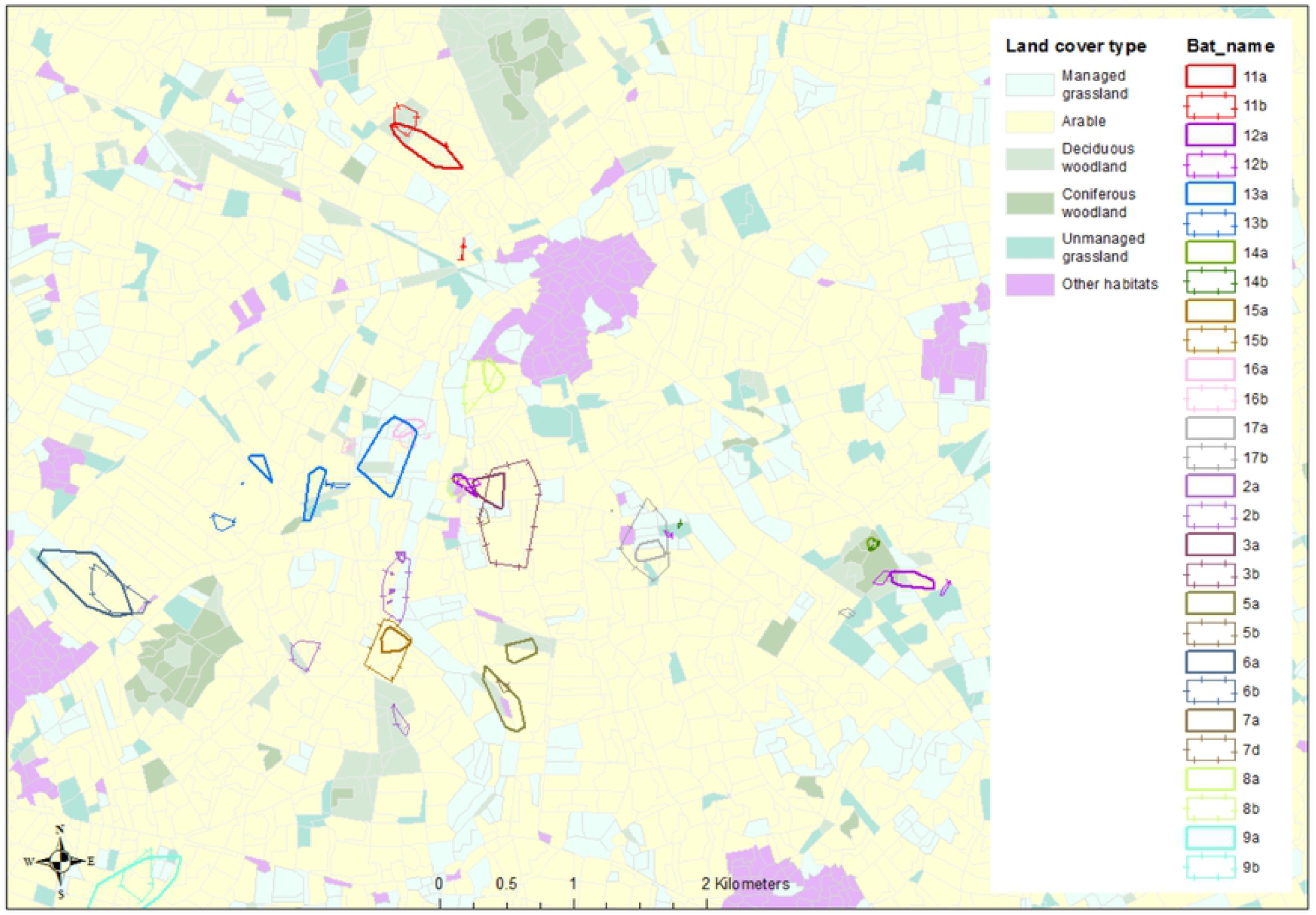
Foraging cores of bats tracked repeatedly at Low Catton (n=14) within the same year (2003) with 2007 Land cover data

**Figure 5.**
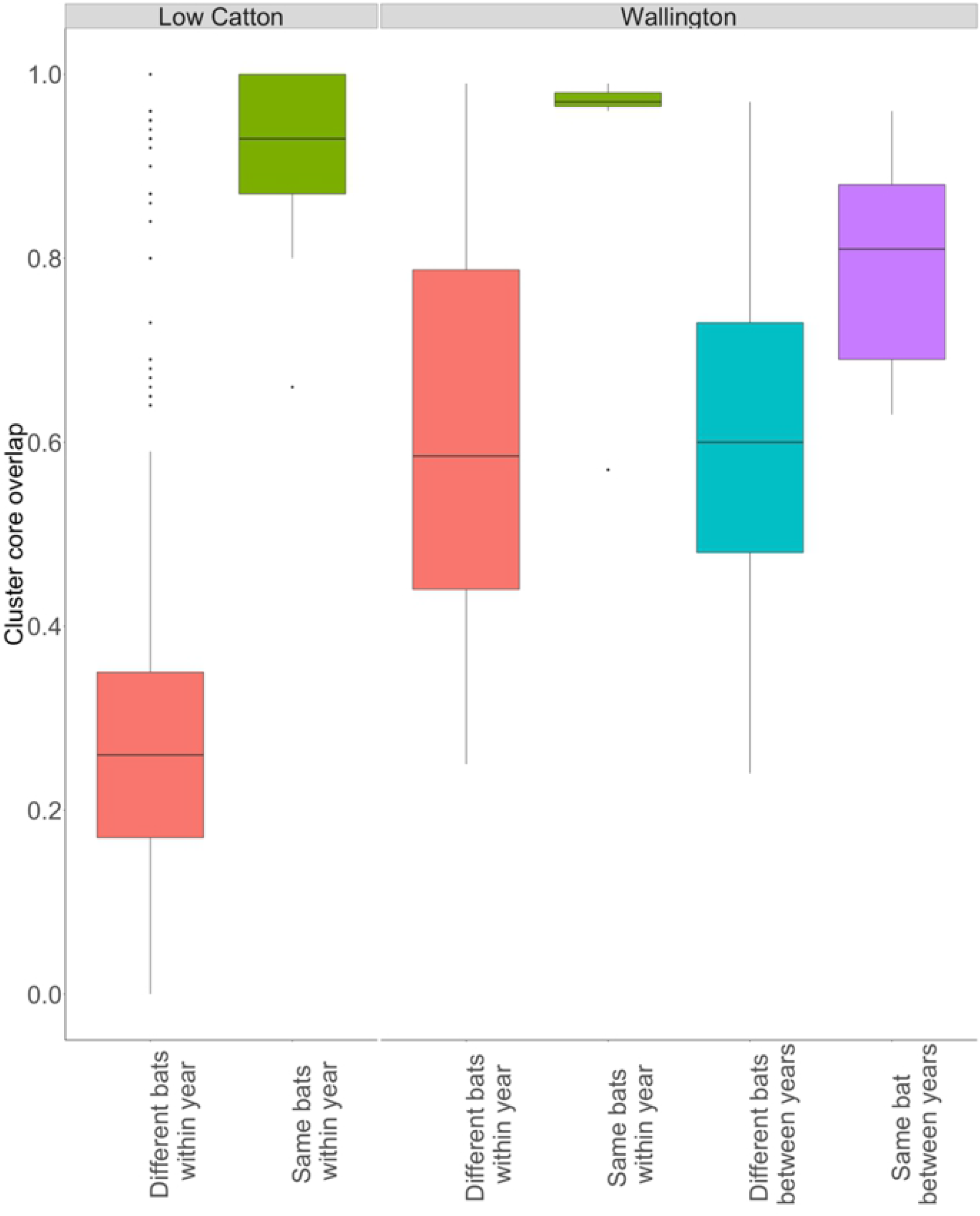
The proportion of overlap of 95% cluster areas for the same bat and different bats within and between years at Wallington and Low Catton. Scores close to 1 indicate 100% overlap of cluster areas whilst scores close to zero indicate independent use of space

There was also a higher degree of spatial overlap for the same bats tracked over different years than for different bats at Wallington (Figure 3, Figure 5; post hoc Tukey tests P=0.02; same bat mean 0.80 ± 0.03 (range 0.63-0.96); different bats mean 0.61 ± 0.01 (range 0.24-0.97)). Individual bats tracked repeatedly also showed significantly more consistency in their foraging strategy (distance to most used foraging core and time taken to travel to most used foraging core) at both sites than pairwise comparisons with other individuals (Table 3).

**Table 3.** Repeatability score (± s.e.) of foraging characteristics of bats tracked at Wallington and Low Catton calculated from ‘rptR’ R package. * indicate repeatability scores significantly higher than random permutations

|  | Wallington | Low Catton |
| --- | --- | --- |
| Distance to most used foraging core | $0.979^* \pm 0.011$ | $0.948^* \pm 0.027$ |
| Time to most used foraging core | $0.698^* \pm 0.132$ | $0.57^* \pm 0.149$ |

### Individual specialisation and habitat use

Overall managed grassland was the most utilised habitat by Natterers at Wallington (11/17 bats had a core are covered by at least 50% grassland) while arable was most utilised at Low Catton (13/17 bats had more than 50% arable in their core foraging area) (Figure 6). However, there was variability in foraging habitat selection by individuals at both sites, for example the proportion of managed grassland within an individual’s core foraging patches at Wallington ranged from 10% to 98%. The population level measure of individual specialisation (WIC/TNW) suggested that, on average, individual bats used a moderate fraction of the total population niche space (0.63 Wallington, 0.69 Low Catton). Monte Carlo analyses of individual versus population niche variation suggested that bats were more specialised than would be expected by chance at both sites (P < 0.001 Wallington, P<0.001 Low Catton) and some individual bats showed very different use of habitat types to others. At Wallington six bats (H1607, H1608, H1609, H1679, H1680, Y2889) used unusual habitats or exploited habitats differently to most of the group (Figure 6) and consequently their PS_i_ values were below the population mean. At Low Catton the majority of individuals had large proportions of arable in their core foraging patches (which was the most dominant habitat type in the area) except for four individuals who showed unique habitat use specialisation (Y2049, Y2106, U8558, U3941) and had large areas of unmanaged grassland, managed grassland, coniferous woodland and a mixture of habitat types respectively (Figure 6).

**Figure 6.**
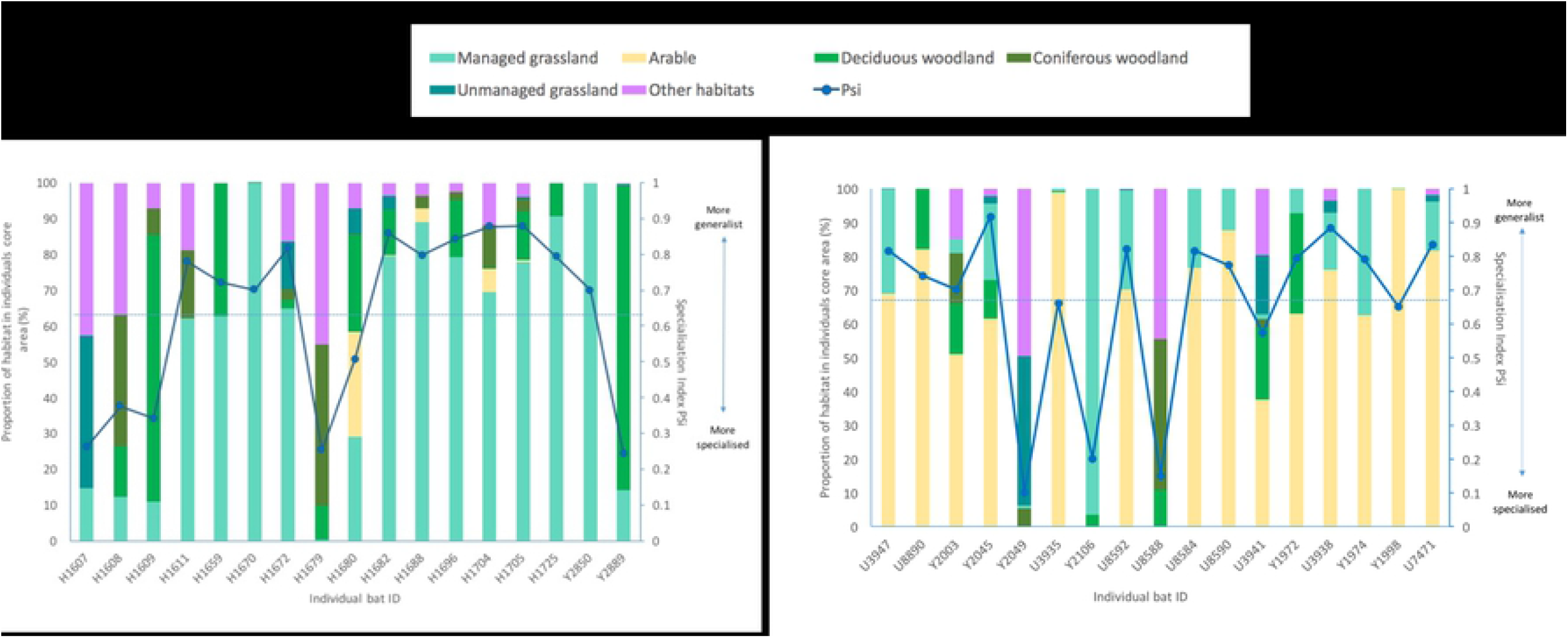
Individual variation in habitat use between bats tracked at Wallington (left) and Low Catton (right). Each individual (x-axis) is represented by a vertical bar, subdivided by the proportion of recordings in each habitat type in relation to the individual’s total cluster core area and the specialization index PSi (blue dots; 0 = more specialized; 1 = more generalist) along with the mean colony Psi (dashed line).

## Discussion

Site fidelity in Natterer’s bats was consistent across a range of intervals (months and years) despite contrasting seasonal contexts. In addition to this the individuals exploited specific foraging locations and showed individual specialisation in their habitat use which is consistent with the behaviour of a territorial species. Bats also exhibited independent choices between the distance to foraging patch and the speed travelled which suggests the theory of optimal foraging for Natterer’s can be rejected.

### Foraging fidelity

Individual foraging fidelity has previously been reported or suggested for a few species of temperate bat (14, 28, 51–53) albeit with variable strength of evidence. More recently, novel methods have extended the suite of bat species that appear to show this behaviour as well as the quality of observation (egert-berg et al 2018). Here, in common with Kerth et al. 2001 and egert-berg, statistically robust quantitative evidence is provided of the re-use of the same locations and habitats by individuals and here this fidelity appears consistent across a range of intervals (months and years) and extends beyond the immediate period of re-tracking defined by the life of currently available radio-transmitters (typically < 10 days for small and medium sized species). Fidelity also appears stable, despite contrasting seasonal contexts (19). This is potentially good news for bat conservation. Firstly, a reliable understanding of the foraging requirements of bats requiring management or conservation is easier to plan for. Once this is understood, the long life of bats suggests that such areas are likely to remain valuable; validating any policy investment in protecting areas of the landscape. Secondly there is considerable value to scientists in knowing that descriptions of individual foraging strategies in adults represent long-lived behaviours, as this helps the integration of foraging choices into spatially explicit studies of the population dynamics of bats in anthropogenic landscapes.

### Territoriality

The behaviour of individuals repeatedly exploiting specific foraging locations whilst also showing individual specialisation in their habitat use is consistent with the behaviour of a territorial species. Here we found that not only do some individuals show striking patch fidelity, but all appear to avoid overlap with other tracked bats which is generally considered to reflect territoriality (53, 54).

This is not the first time that territoriality has been suggested due to apparent foraging site fidelity in individual bats (53–56), though the difficulties in studying individual bats have previously limited authors’ confidence in describing this behaviour. Territoriality has most often been suggested for bat species who have a strong connection with the underlying landscape, and therefore presumably keen to defend a static resource i.e. gleaners or those predating weak flying prey (57, 58). For example, the suggestion of territoriality has often been associated with studies of Daubenton’s bats foraging over still water e.g. (14, 53). For Daubenton’s bat, this speculation is supported by their use of a very specific and easy to map landscape feature, workers’ subsequent confidence in the interpretation of spatial location from radio-tracking and the ability to directly observe foraging behaviour of some individuals. This behaviour includes the repetitive use of space and the presence of social interactions (social calls and chasing) (53). Sociality of temperate bats is becoming widely recognised (17, 18, 58–68) and the identification of social hierarchies within groups is anticipated (69). Whilst social dominance might only be expressed in the choice of roost (69) or the position within the roosting group, it could also be expressed in other key activity bats undertake e.g. foraging; with dominant individuals choosing and maintaining their preferred locations whilst sub-dominant bats may be left with less productive foraging choices. If social dominance does occur, understanding its effects will be important in the management or conservation of bats where pup production is affected by the quality of the foraging resource (70). In the future workers may need to identify and protect the most productive areas of a landscape and potentially distinguish this from the location of preferred habitats, or those habitats used by inefficient or submissive individuals. It should be noted that only approximately half of each social group under study were tracked and there is no evidence of the functional definition of territory defence i.e. observations of antagonistic behaviour between individuals at potential territory sites.

### Behavioural flexibility and individuality

This study agrees with Nachev and York (71) and identifies that individual bats can demonstrate distinct and divergent foraging choices compared to their peers, specifically in their choices of foraging strategy and habitats. In addition, this flexibility in strategy seems to have extended into groups adopting site-specific responses (or traditions) to the contrasting compositions and configurations of the habitats at our two study sites, e.g. the large use of managed grassland at Wallington compared to the use of arable at Low Catton. Further, our demonstration of foraging site fidelity suggests that these individual differences are likely to be long-lived and may represent either differences in personality (71) or tradition in wildlife species, examples of which are now reported for bats (31) as well as other wildlife (30, 33, 72, 73). Alternatively, individual habitat choice may just represent individual preference in a behaviourally flexible species across the broad menu offered by these mixed landscapes. It should be noted that usually, Natterer’s bats are thought to be proficient at both aerial hawking and gleaning (74, 75) and might therefore develop almost unconstrained preferences in prey and varied ways to exploit the foraging options available to them.

## Conclusions

Effective decision-making for conservation and management relies on strong evidence. Here it is shown that individual bats forage at specific core locations, to which they repeatedly return. Whilst it is easy to simply collate the land-cover or habitat types represented within each core into simplistic descriptions of group behaviour, it is clear that individuals differ greatly and show specialisation in their foraging choices, and that for some common habitats, the choice of core location is unique and important and may not be replaceable at another location, even if the habitat appears the same. Thus studies designed to inform conservation and management of temperate bats should attempt to maximize the number of individuals from which movement data is sought, but ensure that data represent a coherent and meaningful measure of behaviour. Further, it is not clear that any of the specific foraging strategies observed at one site (such as commuting style or habitat choices) could transfer to the second, and that our observed behaviours are sensitive to the characters of their landscapes, or the traditions of their communities.

Our finding that the strong foraging site fidelity by Natterer’s bats and our speculation that this may represent some type of territorial behaviour may off-set some of the effort required to collect a single full night of radio-tracking data, as a single full night of data may produce a long-lived and robust description of that bat’s behaviour; justifying the additional effort in field-work hours. It may therefore be a better solution in tracking studies to track many individuals for a single night rather than a few individuals for a longer time frame (76). However it raises the concern that many of the popular analytical methods frequently used by bat workers such as Compositional analysis (77) and selection ratios (78) may be inappropriate. Designing conservation strategies for the rapidly changing environment might then advocate protecting a mosaic of habitats to preserve the habitat specialisms of many individuals rather than choosing a single preferred habitat for a territorial bat which may only suit dominant few individuals.

## Declarations

### Ethics approval and consent to participate

All disturbances at roosts, as well as the capture, handling and marking of bats were carried out under licence from Natural England 2014-6454-SCI-SCI.

### Consent for publication

Not applicable.

### Availability of data and material

The datasets analysed during the current study are available from the corresponding author on reasonable request.

### Authors’ contributions

SM, JA and AM conceived the ideas and designed methodology; SM and JA collected the data; SM and MS analysed data; SM led the writing of the manuscript. All authors contributed critically to the drafts and gave final approval for publication.

## Acknowledgements

We thank Jenni MacKay, Daisy Brickhill, Mark Goddard and Phi Wells for help with the radio-tracking at Low Catton and the National Trust at Wallington for access to their site for catching and radio tracking.

## Additional files

Additional file 1.doc

“Biometric data of bats tracked”

Sex, body mass, reproductive condition, tracking date and number of recordings and locations of bats tracked in this study.

